# Re-profiling of natural inhibitor via combinatorial drug screening: Brefeldin A variant design as an effective antagonist leading to EPAC2 structure modification and antibody design for identification

**DOI:** 10.1101/2021.03.31.437986

**Authors:** Akshay Uttarkar, Vidya Niranjan

## Abstract

Drug discovery can be impactful with re-purposing and combinatorial strategies leading to potential pharmaceutical outputs. Epac2 has been a target of interest for various physiological conditions where suppression is inevitable to achieve desired therapeutic effect. Epac isoforms inhibition is crucial in vascular functions to prevent chronic inflammation leading to hypertension and myocardial infarction. An attempt utilizing brefeldin A, a natural inhibitor was subjected to substitution of 3 side chain groups with 43 fragments via combinatorial strategy. This resulted in generating a library of 79507 brefeldin A variants. High throughput virtual screening yielded 68,043 variants followed by precision docking providing 117 lead like brefeldin A variants. The best docked variant (3-((1*R*,2E,6*R*,10*E*,11a*S*,13*S*,14*A*)-6-(methylsulfonamido)-13-(3-methylureido)-4-oxo-4,6,7,8,9,11a,12,13,14,14a-decahydro-1H-cyclopenta[f][**1**]oxacyclotridecin-**1**-yl)-2,3-dihydro-**1**Himidazol-**1**-ium) has an increased binding efficiency of −10.841 kcal/mol. Simulation studies up to 200ns of complex lead to re-orientation of target tertiary structure resulted in RMSD change of 30.221 Å, suggesting the epac2 structure modification leading to unavailability of RAS-GEF domain and its interaction with Rap1b. A single domain antibody was designed to bind specifically to re-structured epac2 for potential identification over the native target structure. The resulting Brefeldin variant can be potentially labelled as a most effective antagonist against epac2 which induces theoretically irreversible structural re-conformation. This study also provides a robust *in-silico* workflow for searching of chemical space, generating and screening of combination libraries and the efficient utilization of known inhibitor.

## Introduction

Exchange protein directly activated by cAMP2 (Epac2) is a protein coded by RAPGEF4. It interacts with Rap1b via Cyclic adenosine monophosphate (cAMP) mediated mechanism. cAMP binds to the cNB-DB domain inducing a conformational change to epac2 structure. This change enables the activation of catalytic region for interaction with Rap1b. [1]

Epac2/Rap1b pathway has a significant physiological role in vascular isoforms. The effect of cAMP in preventing inflammation of vascular tissues particularly by interleukin (IL-6) propagation has well studied [2], Fluorescent cAMP analogue has provided insights into Epac2 selective inhibitors. The work focusses on competitive binding of 14,400 molecules in NDB-cAMP domain [3]. Amidst, all the efforts of targeted inhibition of epac2 we find many reports suggesting denaturing of target proteins. Such reports raise grave questions regarding non-specific binding leading to denaturation [4], This brings us to providing a specific compound from a known derivative which causes structural re-confirmation and prevent denaturation. Overall, the status of antagonist to epac2 issues need to re-addressed by an initiative approach by reprofiling of known inhibitors to achieve desired functionality.

To achieve the particular re-purposing of drugs, a robust chemical fingerprinting and combinatorial screening protocols have to be standardised. This prevents non-specific and cross-interactions between proteins of same family. For anticancer therapy, to avoid cross talks between proteins in signalling pathway a combination drug research lead to increased efficiency of therapeutics [5]. Generation and screening of combination libraries is crucial to avoid generation of false positives, leading to skewed results. Understanding of excluded volumes via pharmacophore studies provides in depth insight on the chemical space occupied and impact of the same on binding efficiency.

In recent studies, the antagonists have shown lack of confidence in results for *In-silico* and *In-vitro* of epac2 signalling pathway [6]. The study carried out is an attempt to deliver a workflow to achieve stability in in-silico study.

In the current study, Brefeldin A has been our choice of antagonist which can be re-profiled. It is a small hydrophobic compound having impactful effects on mammalian cells produced by *Penicillium brefeldianum*. It is a potential inhibitor against activation of ADP-ribosylation factors and small G protein of Arf family (7).

All the docking experiments carried out should be followed by a simulation study for a minimum of 100ns to confirm stability of binding molecule and the structural variations caused to the target molecule due to stable interactions.

The combination of above-mentioned approach appeals to finding a better antagonist that provides solution to current problem of target degradation. The major focus of the study is the re-modification of the target structure and developing a single domain antibody (sdAb) for detection of the target-ligand complex.

## Methods

### Domain and structural analysis

The protein structure for Epac2 is available in Protein data bank, with following id’s 4FHZ, 3CF6, 4MH0 and 1O7F. For the current study 4FHZ (*Mus musculus*) and 3CF6 (*Homo sapeins*) have been selected based on the coverage of amino acid sequence and known protein interaction in Rap1b. Similarity, Identity and structure superimposition was carried out with 4MH0 and 1O7F for validation.

Domain analysis for epac2 was conducted using InterPro and Conserved domain database of NCBI. A schematic of complete workflow is shown in (Fig 6).

**Fig 1.**
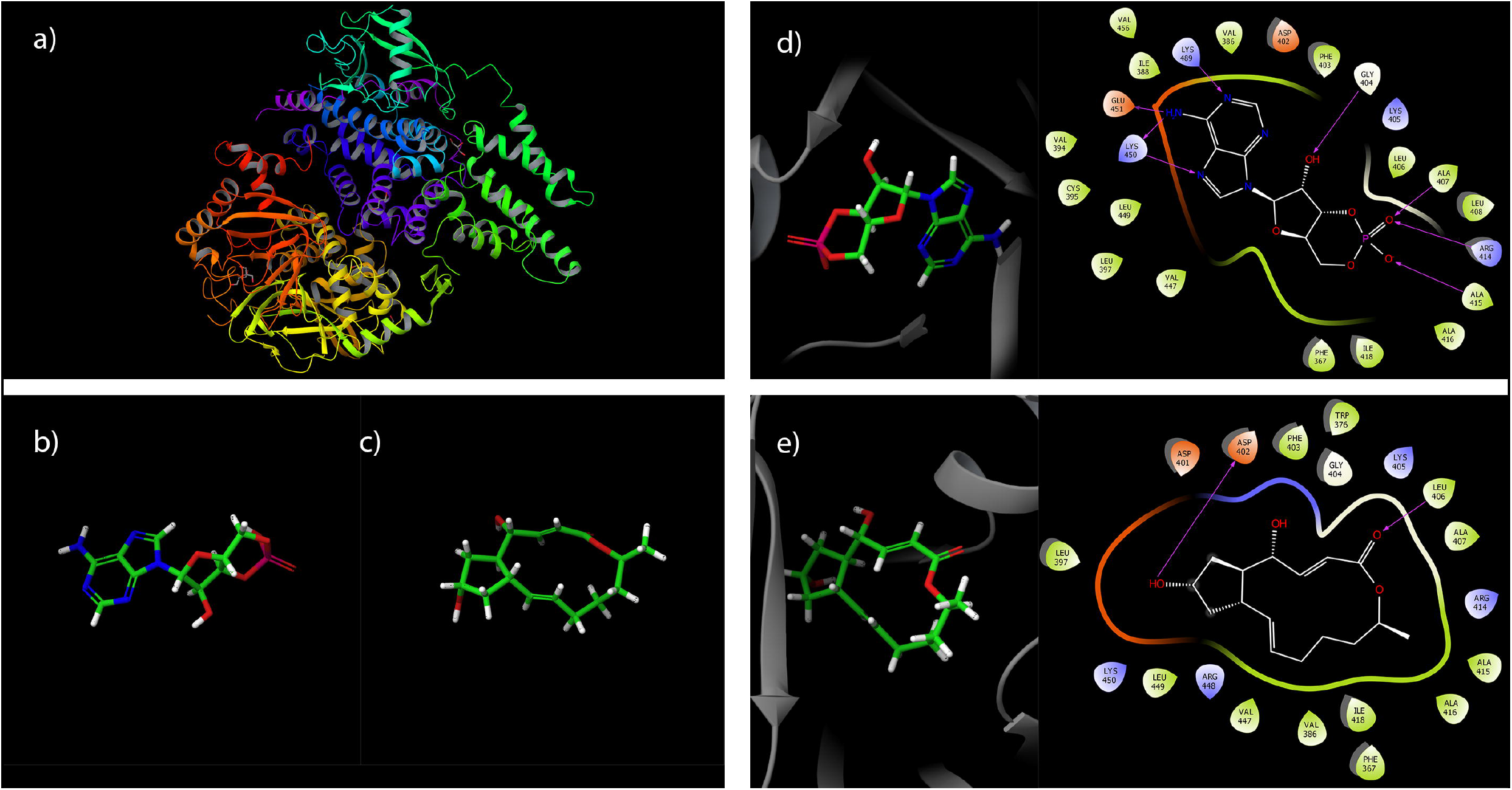
Structural information of target and compounds, along with interactions. (a) Epac2 structure colour coded highlighting structural and catalytic domains (b) Structure of cAMP (c) Structure of Brefeldin A (d) Interactions of cAMP in the binding pocket, along with a 2D interaction diagram exhibiting the interaction with epac2. (e) Interactions of BrefA in the binding pocket, along with a 2D interaction diagram exhibiting the interaction with epac2. The above images were developed using Maestro 2019-3 (https://www.schrodinger.com/maestro), available free for academic usage.

**Fig 2.**
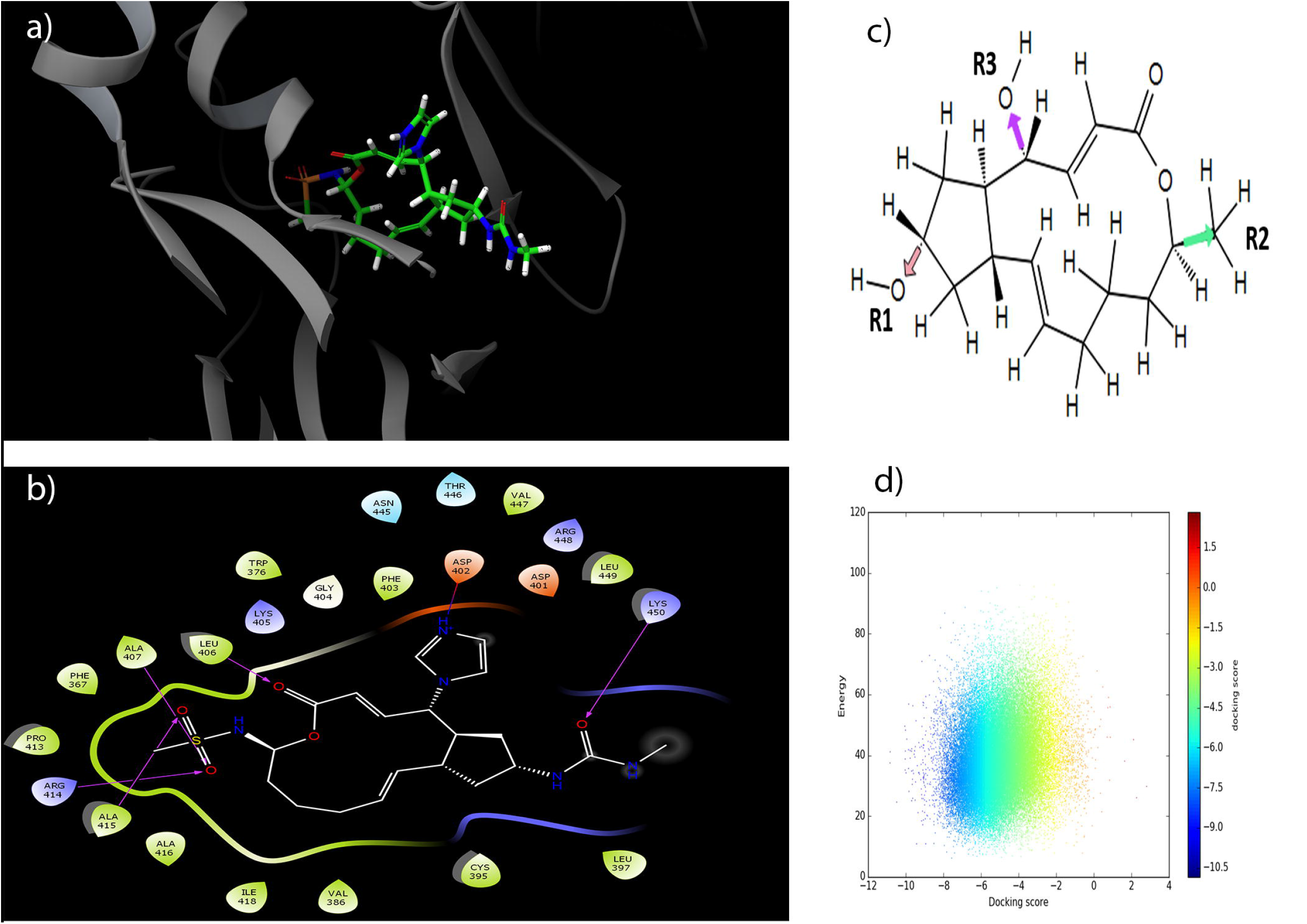
Selection of Brefeldin A side atoms, combinatorial screening and docking plot along with Brefeldin A variant interactions. (a) BrefA best variant (BrefA1) in the binding pocket of EPAC2 (b) BrefA1, 2D interaction diagram exhibiting the interaction with epac2 (c) 2D diagram of Brefeldin A showing core and side chain atoms Rl, R2 and R3 were replaced with 43 selected fragments in permutation and created a combinatorial library of 79507 BrefA variants, (d) Docking score vs Energy plot for HTVS, SP and XP. Representation via gradient colour coding of docking score to show selection of variants via better and efficient docking score. The above images were developed using Maestro 2019-3 (https://www.schrodinger.com/maestro), available free for academic usage

**Fig 3.**
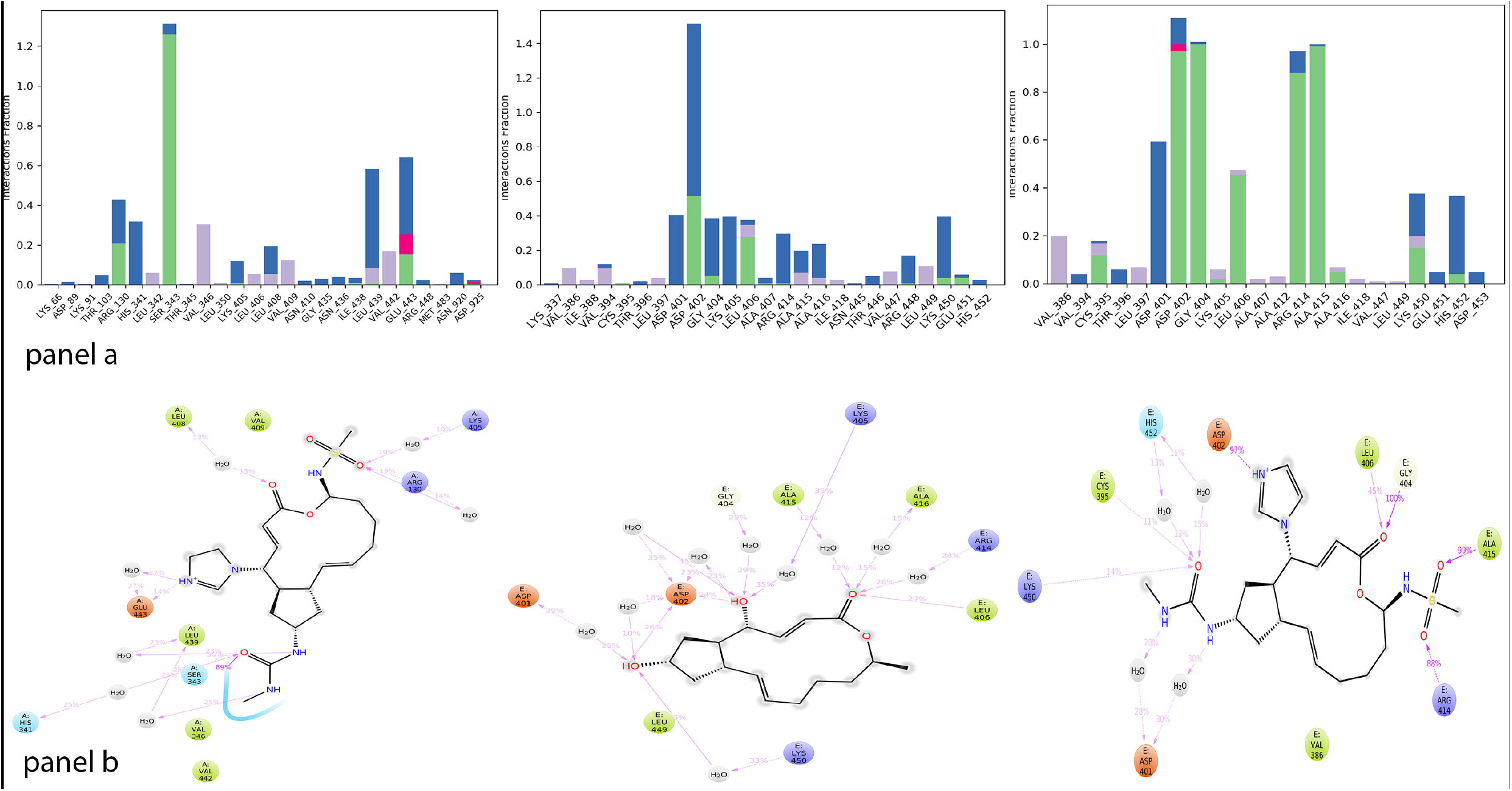
2D Interactions and protein-ligand contact histogram for 200ns. (Panel a) from left consists of epac2-cAMP, epac2-brefA and epac2-brefA1 complexes subjected to 200ns simulation. The interaction of ligands with residues exhibiting over the course of simulation are recorded. The histogram (Interaction fraction vs Residues) suggests the percentage of specific type of interaction of an amino acid with ligand. Green suggests hydrogen bonding, Blue for water bridges and pink for ionic interactions. (panel b) from left consists of 2D interaction for epac2-cAMP, epac2-brefA and epac2-brefA1 complexes respectively. The plot suggests respective interactions at the end of simulation has been shown along with percentage contribution of each interacting residue with the compound. The above images were developed using Desmond 2019-2 (https://www.deshawresearch.com/resources_desmond.html) available free for academic and non-commercial usage.

**Fig 4.**
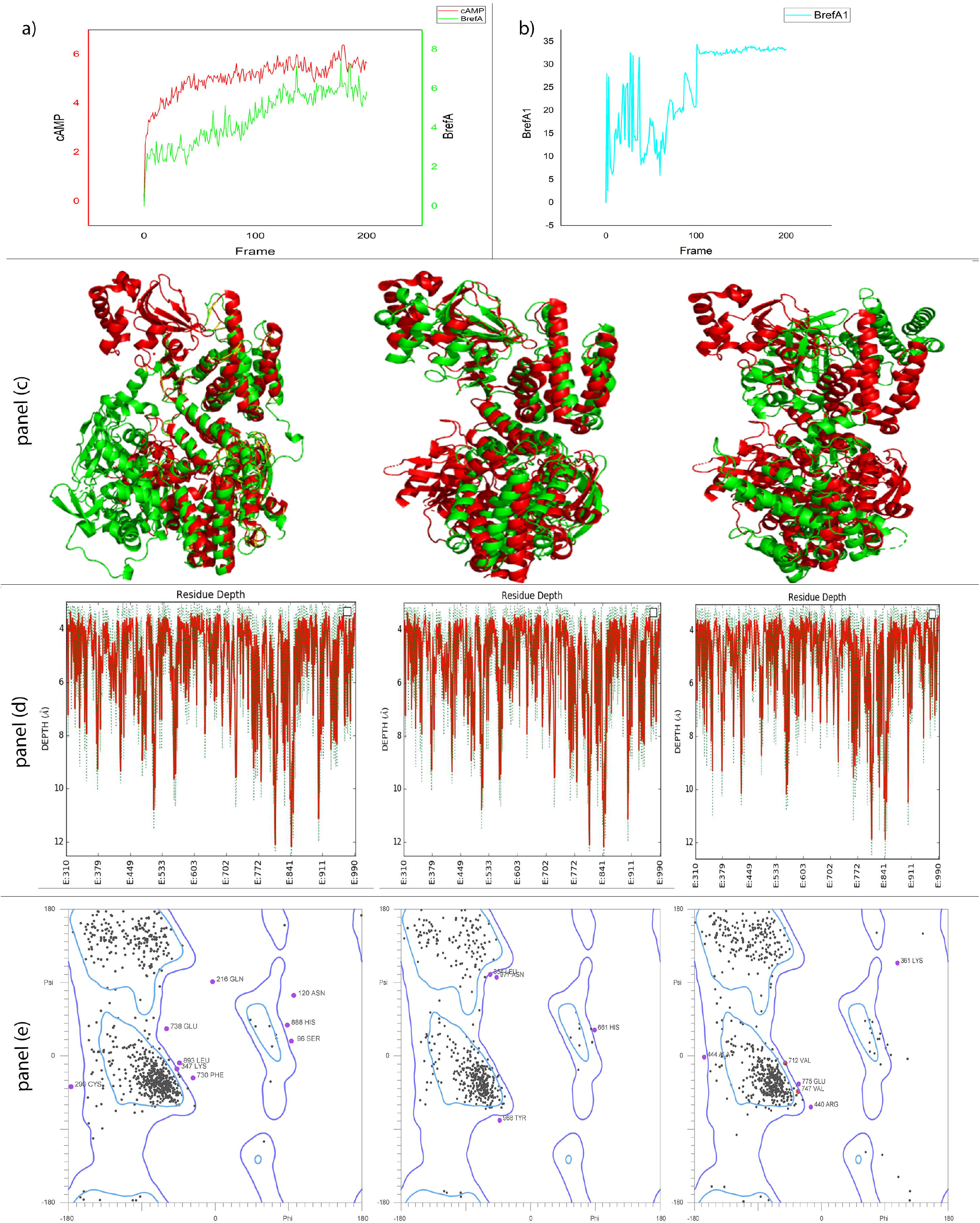
Analysis of structural conformations exhibited by epac2 on interaction with cAMP, BrefA and BrefA1. (a) A dual axis plot of RMSD (Å) vs time frame (ns) plots for epac2-cAMP and epac2-BrefA show a similar change in deviation. Epac2-cAMP in red and epac2-BrefA in green rescaled to fit and highlight the change in RMSD. (b)Epac2-BrefA1 in cyan exhibit the RMSD change versus time. The above figures were plotted using R studio 1.1.456 (https://rstudio.com/products/rstudio/download/) and ggplot libraries (https://rstudio.com/products/rpackages/), free open source licensing available. (Panel c) From left, Epac2-cAMP, Epac2-BrefA and EPAC2-BrefA complexes respectively were subjected to superimposition over the reference structure. Green corresponds to final structures after simulation over the reference structure (3CF6) in Red. The above images were developed using pymol version 2.3.2 (https://pymol.org/2/), available on trial version. (Panel d) From left, Epac2-cAMP, Epac2-BrefA and EPAC2-BrefA complexes respectively the residue depth is calculated and the residue versus depth in Å from the centre of the protein in plotted. The peaks in red refer to surface residues and gradient colouring to green for buried residues. The above images were obtained from DEPTH server (http://cospi.iiserpune.ac.in/depth/htdocs/index.html) (Panel e) From left, Epac2-cAMP, Epac2-BrefA and EPAC2-BrefA complexes respectively the Ramachandran plot of the protein structures after simulation to check structural stability. The images were plotted using Molprobity (http://molprobity.biochem.duke.edu/)

**Fig 5.**
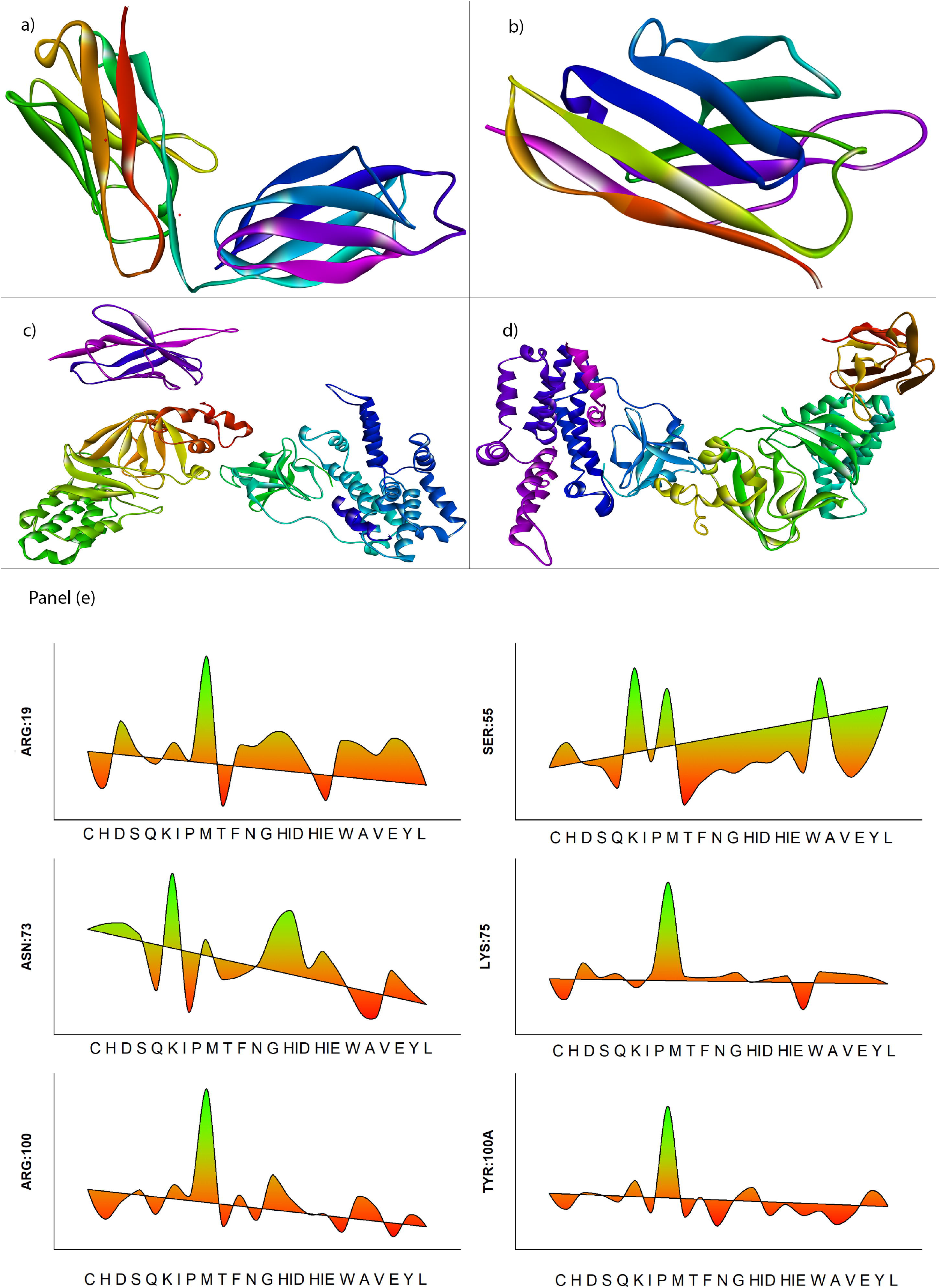
Antibody design and Antigen-antibody docking via mutagenesis. (a) The template of 1DEE_B used for modelling the framework and loops of antibody (b) Modelled antibody representing H1, H1 and H3 regions highlighted in red, green and purple respectively, (c) Native Epac2 interaction with single domain antibody (d) Epac2 interaction with mutated single domain antibody. The above images were developed using Maestro 2019-3 (https://www.schrodinger.com/maestro), available free for academic usage. (Panel e) A graphical representation of the most interactive antibody residues (x axis) subjected to mutagenesis plotted against alternative amino acids (y axis) with the solvability score. The antibody was substituted with residues with highest solvability score. The δ solvability scores can be obtained from table 3. The above stack plot was plotted using R studio 1.1.456 (https://rstudio.com/products/rstudio/download/) and ggplot libraries (https://rstudio.com/products/rpackages/), free open source licensing available.

**Fig 6.**
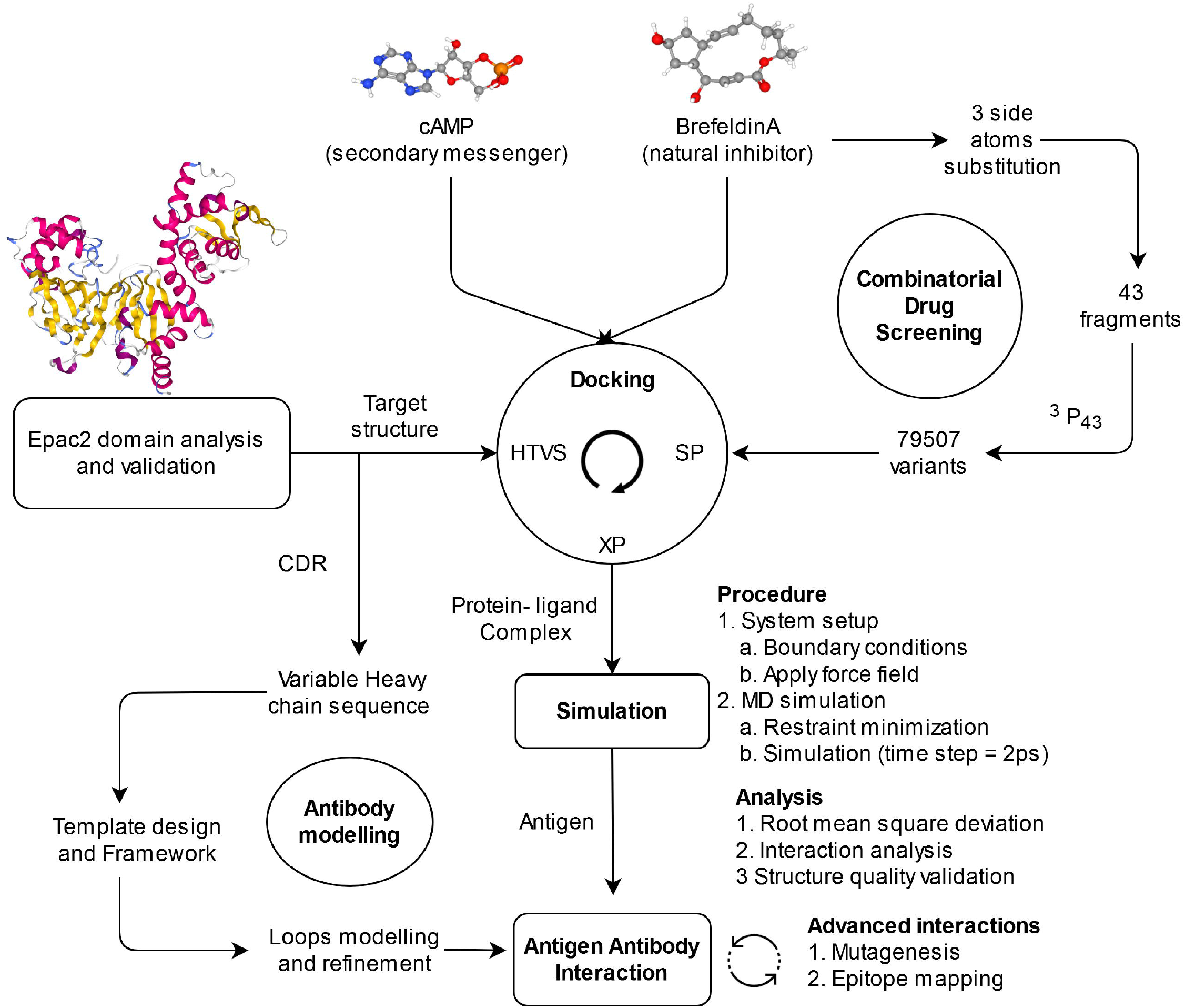
Schematic representation of methodology. The overall objective of the method designed in to re-profile the Brefeldin A (natural inhibitor) into an antagonist and achieve desired conformational changes to EPAC2 structure. Docking studies were performed for cAMP and BrefA. 3 side chain groups of BrefA were further subjected to substitution with 43 fragments, leading to a database of 79507 variants of BrefA. High throughput virtual screening (HTVS) of brefA variants, followed by Standard precision (SP) and Extra precision (XP) docking with EPAC2. The protein complexes EPAC2-cAMP, EPAC2-BrefA and EPAC2-BrefA1 (best docked variant) were subjected to simulation studies for 200ns. Laterally, the complementary determining regions (CDR) for EPAC2 were targeted for developing a single domain antibody (sdAB), which was refined by epitope mapping and mutagenesis. The aim was to have a sdAB, exhibiting preferential binding towards EPAC2-brefA1 complex over other complexes and native structures. The above schematic representation was created using Draw.io (https://www.draw.io/)

### Combinatorial drug screening

The Small molecule Discovery suite 2019-3 was used for Docking, R-group enumeration and virtual screening. Various tools like Protein Preparation Wizard [8] Epik (9), Impact (10), LigPrep [11], SiteMap [12], Receptor grid generation and Glide [13] were utilized.

The protein was subjected to pre-processing for defining residues, HET atoms and verify valency. Under structure refinement, hydrogen bond assignment was carried out for neutral pH. Structure minimization for less than 0.30 Å was conducted with OPLS3e defined force field. The top ranked potential binding sites were selected with restricted hydrophobicity. The in the present case, the cAMP2 binding domain was selected as a binding pocket which was evaluated as top binding site. A receptor grid was defined for docking of BrefeldinA and cAMP with the selected binding pocket with constraints and predefined excluded volumes. LipPrep was used to generate 3D structures of ligand molecules with 32 conformers for each respectively. An extra precision (XP) ligand docking was performed using Glide, with a scaling factor of 0.8 and partial cut-off of 0.15. Epik state penalties were added to the docking score.

The Enumeration and Ideation tool [14] was used for preparation of custom R-Group libraries. A custom R groups of 43 fragments was created and was utilized was substitution with 3 side chain atoms of BrefeldinA. A total of 79,507 variants of Brefeldin A were generated. This generated library was subjected to LigPrep and stored as a database for further screening.

A virtual screening workflow was created. The ligand input files were selected from the database created. A pre-filter was defined to follow Lipinski’s rule using QikProp [15] properties. Epac2 protein structure was selected as receptor with defined binding site developed using Sitemap and receptor grid. A docking workflow for High throughput virtual screening [HTVS] to dock flexibly, perform post-docking minimization and generates one pose per compound state. All states are retained after docking. The top 10% of compounds are carried forward for Standard precision (SP) docking. The parameters remain the same as selected for HTVS. The top 10% are carried forward for Extra precision (XP) docking. For XP, except for generating 3 poses for compound state, remainder parameters are constant The final lead like compounds are subjected to post processing with prime MM-GBSA for accuracy in binding efficiency. The best docked compound was merged with the target protein structure and the complex was exported as .pdb file from the workspace navigator.

The 2D interaction plot was generated from ligand interaction viewer in Maestro [16]. The 2D plots for docking score vs energy was generated using Plot viewer tab in tables section of Maestro.

### ADME studies

ADME studies were performed on SwissADME web server [17] by importing the compound structure, conversion to SMILES and prediction.

### Simulation studies

Desmond [18] was used for performing simulations. The interaction complex was subjected to protein pre-processing and H-bond assignment with similar parameters as mentioned earlier. Simulation system was built utilizing the system builder. The solvent model selected was TIP3P, boundary conditions defined by orthorhombic box with minimized volume encapsulating the complex. The force field applied is OPLS3e. System was neutralized by adding Cl- or Na+ ions based on the system total charge.

To perform molecular dynamics simulation, the system was imported from the workspace. Simulation time was 200ns, with trajectory recording interval set to 1ns (the optimised interval time helps in clear visualization and appropriate storage space). Ensemble class was set to NPT (defines the thermodynamic parameters of constant pressure and temperature, a variable volume allows the protein to undergo conformational change) and modelled relaxed for 10 ns before performing simulation (a relaxation protocol avoids increase in atoms velocity due to increase in temperature). Time step was set to 2 femto-seconds (fs) and temperature of 300K. A cut-off short range method with a radius of 9.0 Å (This optimized value avoids overlapping of atoms). No restraints were pre-defined.

Simulation has a temperature increase of 10K per time step after solvation of binding pocket.

Simulation interaction diagram tool was used for analysing the simulation results for change in RMSD and protein-ligand contacts. The surface depth analysis was carried out using the DEPTH server [19]. Protein structure superimposition calculation and visualisation was carried out by Pymol [20].

The stability of protein structures after completion of simulation course was evaluated by Ramachandran plot utilizing Mol-probity web server [21],

### Antibody design

The Biologies suite [22] provides tools for antibody design, antigenantibody interaction and mutagenesis. The human heavy variable chain sequence (A2NXP8) was selected as sequence for antibody design. Under antibody prediction tool, for single domain antibody the selected sequence was provided as input. A homology search was conducted for shortlisting of antibody template structure. The top most template was selected as antibody framework. The H1, H2 and H3 loops of the CDR regions were modelled and the best model was reported back.

### Protein Protein docking

Protein protein docking tool was utilized with antibody mode masking the non-CDR regions. The antigen and antibody were selected for docking. Number of rotations to probe was set to 70000 and maximum poses to return set to 30.

No attractions-repulsions and distance restraints were defined. The protein interaction visualization was used for epitope mapping and interaction calculation.

The protein-protein docked complex was subjected to antibody humanization-residue mutation to improve stability of protein-protein interaction. The critical residues were subjected of substitution with reminder of amino acids to calculate affinity, SASA, pKa and complementarity.

The Residue mutation viewer was used to plot the change in delta stability results. The amino acids substitution was carried out in sequence viewer. The protein-protein docking was carried out for mutated antibody and antigen. The docking parameters and visualization was carried out as mentioned earlier.

## Results and Discussion

### Domain and Structural analysis

The domain analysis of the epac2 was carried out (Fig 1a). The cAMP1 ranges from 43-166 amino acid (aa), DEP ranges from 198-273 aa, cAMP2 ranges from 338-458 aa, N-terminal RAS-GEF ranges from 480-616 aa and RAS-GEF ranges from 757-934. cAMP domain is further classified into PBC ranging from 403-417 aa and hinge ranging from 433-438 aa. The cAMP binding at the PBC region triggers the hinge mechanism providing RAS-GEF domain an interacting surface for Rap1b.

Three structures (4F7Z, 4MH0 and 1O7F) of epac2 are available in PDB and 4F7Z was selected. Similarity, identity and structure superimposition was carried out with PDB id’s 4MH0 and 1O7F for validation purposes of protein structure taken into consideration. 4F7Z had similarity of 98.36% and 91.17% with 4MH0 and 1O7F respectively. A structural superimposition revealed a RMSD variation of 1.21 Å and 1.68 Å with 4MH0 and 1O7F respectively. Based on the observations, we can conclude that 4F7Z has structural similarity to 4MH0 and 1O7F. Since 4F7Z has better sequence coverage it has been selected for docking studies. The sequence coverage can be visualized in (Supplementary Fig 1)

### Combinatorial drug screening

#### Comparison of cAMP and Brefeldin A interaction with Epac2

As discussed in the previous section, regarding the hinge mechanism both cAMP (Fig 1b) and Brefeldin A (BrefA) (Fig 1c) have shown highest binding efficiency in the cAMP2 domain. We have performed extra precision docking of cAMP and BrefA with EPAC2. cAMP has a binding energy of – 10.692 kcal/mol whereas BrefA has a binding energy of −6.248 kcal/mol. cAMP forms conventional hydrogen bonds with D402, G406, A407, A415, K450 and has an attractive charge interaction with R414 (Fig 1d). Whereas BrefA exhibits hydrogen bonding with D402 and L406 (Fig 1e). It can be concluded that cAMP undergoes efficient binding due to more significant interactions in the binding pocket. A higher binding energy and stronger interactions promotes the hinge mechanism with cAMP. BrefA is a competitive binding molecule aimed to avoid cAMP binding.

#### Generation of combinatorial library

Three sites on the Brefeldin A molecule were selected by avoiding core atoms for replacing it with 43 side chain fragments. The sites selected were (R1) C9 linked with - OH (O1 and H31) with bond length of 1.423 Å (R2) C10 linked with −OH (O2 and H33) with a bond length of 1.428 Å and (R3) C18 linked −CH3 (C20, H42, H43, H44) with a bond length of 1.523 Å (Fig 2c). A library of 43 fragments was built which will be in permutation be replaced with each of the selected sites. This resulted in a library of 79507 BrefA variants (Table 1) with a common core similar to of BrefA.

**Table 1.** List of side chain fragments used in Brefeldin A variants library.

| SI<br>no | Fragment | SI<br>No | Fragment | SI<br>no | Fragment |
| --- | --- | --- | --- | --- | --- |
| 1 | methane | 16 | Ammonium | 31 | N,N-dimethylmethanesulfonamide |
| 2 | propane | 17 | methylammonium | 32 | benzenesulfonamide |
| 3 | tert-butane | 18 | dimethylammonium | 33 | benzenesulfonamide 2 |
| 4 | trifluoromethane | 19 | hydrogen cyanide | 34 | N-methylbenzenesulfonamide |
| 5 | cyclohexane | 20 | Water | 35 | N-methylurea |
| 6 | benzene | 21 | Acetamide | 36 | N-phenylurea |
| 7 | pyridine 1 | 22 | acetamide 2 | 37 | morpholine |
| 8 | pyridine 2 | 23 | acetamide 3 | 38 | morpholinium |
| 9 | pyridine 3 | 24 | N-methylacetamide | 39 | piperidinium |
| 10 | imidazole 1 | 25 | N-methylacetamide<br>2 | 40 | formate 1 |
| 11 | imidazole 2 | 26 | N,N-dimethylacetamide | 41 | tetrazolate |
| 12 | imidazole 3 | 27 | Acetanilide | 42 | N-methylmethanesulfonamide |
| 13 | imidazolium 1 | 28 | methanesulfonamide | 43 | N-methylmethanesulfonamide 2 |
| 14 | imidazolium 2 | 29 | methanesulfonamide<br>2 |  |  |
| 15 | imidazolium 3 | 30 | methanesulfonamide<br>3 |  |  |

#### Virtual screening of Combinatorial library

High throughput virtual screening (HTVS) of library compounds resulted in docking of 68,043 variants. The remainder of 11,805 variants could not be bound to target. The reason behind omission is compound surface area is larger than the size of binding pocket. The top 10% of the HTVS docked molecules based on binding efficiency i.e 6,804 compounds were subjected to standard precision (SP) docking. The top 10% of the SP docked molecules i.e 680 molecules were subjected to Extra precision (XP) docking with 10 best poses to be reported. The final set of 117 variants were shortlisted. The threshold of selecting 117 lead like variants was their binding efficiency was > −6.248 kcal/mol (binding energy of Brefeldin A). These variants were subjected to molecular mechanics combined with poisson-boltzmann surface area (MM-GBSA) to calculate the accurate binding energy. The lead molecules docking information is provided in (Supp table 1). The docking score and energy of the docked molecules are shown in (Fig 2d). The best docked variant of Brefeldin A is Brefeldin A; [N-methylurea 1];[imidazolium 2]; [from methanesulfonamide 3]. The IUPAC name of the obtained variant is (3-((1R,2E,6R,10E,11aS,13S,14aR)-6-(methylsulfonamido)-13-(3-methylureido)-4-oxo-4,6,7,8,9,11a,12,13,14,14a-decahydro-1Hcyclopenta[f][1] oxacyclotridecin-1-yl)-2,3-dihydro-1Himidazol-1-ium). It has an increased binding efficiency of −10.841 kcal/mol. The best docked variant will be now onwards referred to as BrefA1.

The details of fragments replaced is R1 replaced with methylurea 1, R2 replaced with imidazolium 2 and R3 replaced with methanesulfonamide 3. The docked BrefA1 in the binding pocket and 2D interaction with EPAC2 is shown in (Fig 2a and Fig 2b).

## Absorption Distribution Metabolism Excretion (ADME) studies of Brefeldin A1

In (Table 2), a detailed analysis of five components like water solubility, pharmacokinetics, physicochemical, drug likeness, lipophilicity and medical chemistry of Bref A1 has been reported.

**Table 2.** Featuring various properties regarding ADME of Brefeldin A1.

| Water Solubility |  | Pharmacokinetics |  |
| --- | --- | --- | --- |
| Log S (ESOL) | -3.37 | GI absorption | Low |
| Solubility | 2.00e-01 mg/ml; 4.27e-04 mol/l | BBB permeant | No |
| Class | Soluble | P-gp substrate | Yes |
| Log S (Ali) | -4.21 | CYP1A2 inhibitor | No |
| Solubility | 2.88e-02 mg/ml ; 6.16e-05 mol/l | CYP2C19 inhibitor | No |
| Class | Moderately soluble | CYP2C9 inhibitor | No |
| Log S (SILICOS-IT) | -2 | CYP2D6 inhibitor | No |
| Solubility | 4.65e+00 mg/ml ; 9.93e-03 mol/l | CYP3A4 inhibitor | No |
| Class | Soluble | Log Kp (skin permeation) | -8.01 cm/s |
| Drug likeness |  | Physicochemical Properties |  |
| Lipinski | Yes; 0 violation | Formula | C21H34N5O5S |
| Ghose | Yes | Molecular weight | 468.59 g/mol |
| Veber | No; 1 violation: TPSA>140 | Num. heavy atoms | 32 |
| Egan | No; 1 violation: TPSA>131.6 | Num. arom. heavy atoms | 0 |
| Muegge | Yes | Fraction Csp3 | 0.62 |
| Bioavailability Score | 0.55 | Num. rotatable bonds | 6 |
| Medicinal Chemistry |  | Num. H-bond acceptors | 7 |
| Brenk | 1 alert: isolated_alkene | Num. H-bond donors | 4 |
| Leadlikeness | No; 1 violation: MW>350 | Molar Refractivity | 129.03 |
| Synthetic accessibility | 6.05 | TPSA | 141.83 Å<sup>2</sup> |
| **Lipophilicity** |  |  |  |
| Log Po/w (iLOGP) | 1.82 | Log Po/w (MLOGP) | -2.93 |
| Log Po/w (XLOGP3) | 1.62 | Log Po/w (SILICOS-IT) | -2.22 |
| Log Po/w (WLOGP) | 0.02 | Consensus Log Po/w | -0.34 |

Bref A1 is found to be soluble in water, with Log S values suggesting the same, druglikeness study satisfies all the parameters including the Lipinski’s rule, Ghose filter and a bioavailability score of 0.55. The topological polar surface area (TPSA) is 141.83 Å^2^, an expected threshold is TPSA < 140 Å^2^. The permeability with cell surface is still feasible but with low rate of absorption.

Blood brain barrier permeability is not possible with TPSA score >90 Å^2^. The synthetic accessibility score is 6.05 which makes the compound moderately feasible to synthesize.

XLogP3 value of 1.62 is within the range of −0.7 < XLogP3 > +5 ensuring favourable dissolution in non-polar solvents.

## Molecular Dynamics Simulation studies

Simulation study was performed for three protein-ligand complexes namely epac2-cAMP, epac2-BrefA and epac2-BrefA1 (Bref A best docked variant). Each of the complex was subjected to 200ns simulation in a neutral orthorhombic environment.

### Simulation interaction analysis

An in-depth analysis of P-L contacts was conducted to confirm the stability of ligand binding efficiency. Throughout the length of the simulation, the ligands were engaged in stable interactions with target residues. The interactions of epac2-cAMP remain consistent with G404, A407, R414, A415, K450, K489. K450 exhibits hydrogen bond interaction with N1 and N2 of adenosine ring, throughout the course of simulation. Epac2-BrefA interactions are D402, L406. Both the interactions exhibit hydrogen bonds significantly mediated via water bridges. Epac2-BrefA1 have interactions with D402, L406, A 407 R414, A415 and K450. All the residues have exhibited stable hydrogen interaction, with D402 forming salt bridge in the simulation course. Consideration of hydrogen-bonding properties in drug design is important because of their strong influence on drug specificity, metabolization and adsorption. The interaction diagram (Fig 3 panel b) suggests stronger interaction in epac2-brefA1 complex with higher number of hydrogen bonds.

In (Fig 3 panel a), protein ligand interaction is summarized and stacked as bar plots suggesting “a particular type of interaction” over a total period of simulation. A score of 0.5 indicates that for 50% of total simulation time the interaction existed. A score >1.0 can exist and suggests that more than one atom is involved in interaction. The stable complex of epac2-brefA1 has D402 forming hydrogen bond, ionic bond mediated via water bridge. A G404, R414 and A415 have shown strong hydrogen bonding with minimalistic mediation of water bridge. These stable interactions have greater impact on the stable conformational change induced onto EPAC2.

#### Root Mean Square Deviation (RMSD) of protein structure

Monitoring RMSD calculation for C-α atoms of all residues over the course of the simulation gives in-sights into the structural confirmation changes undergoing. Changes of the order 1-3 Å are perfectly acceptable for small and globular protein. Larger the change, higher will be the protein structure evolving the course of simulation. All the three complexes were subjected to simulation of 200ns to study the changes induced by ligand interactions.

Both epac2-cAMP and epac2-BrefA have a similar RMSD change of 5-6 Å (Fig 4a). The epac2-cAMP on contrary exhibits an initial shift which results in hinge mechanism and availability of RAS-GEF domain for interaction. On the contrary, epac2-BrefA1 showed a lot of variation. In (Fig 4b), it can be clearly visualized that from 1 to 40 ns time frame the fluctuations were ranging from 1 to 32 A. From 40 to 100ns time frame, the protein structure evolved by a stabilized increasing trend in RMSD values. After 105 ns, the fluctuations in RMSD stabilized suggesting the protein structure attaining stable conformation.

The changes in the tertiary structure of epac2 can be compared by superimposition of simulated structure with reference structure (PDB id - 3cf6). In (Fig 4 panel c), a compare and contrast view on the change in structure is provided. Epac2-brefA superimposition is almost in alignment, as which is in terms with literature evidence. The epac2-cAMP on superimposition provides a clear picture on the conformational change induced, hinge movement and availability of RAS-GEF domain for further interaction with Rap1b. Epac2-brefA1 interaction is in no alignment with the reference structure validating the change epac2 conformation.

The changes in the protein structure at the end of simulation has been captured by calculating the residue depth for each of the complex. In (Fig 4 panel d), the distance of each residue from the centre of the structure is calculated via distance (Å). Based on this calculation, the nature of a residue whether buried or exposed to surface is plotted. This change in peaks suggest change in residue position and structure. This further validates the change in RMSD and superimposition results.

Conclusion drawn is binding of BrefA1 in cAMP2 domain induces a complete re-conformation of EPAC2 structure. This change can be justified purely due to increase in binding efficiency of BrefA1 relative to cAMP. The steric effect in protein is also due substituted side chain atoms of BrefA1 and its interaction with cAMP2 domain.

The brefA1 induced changes are not leading to any breakage in protein structure and that the stability is attained in course of simulation.

### Protein stability evaluation

Each of the docked complex at the end of simulation was subjected to protein structure quality evaluation. Ramachandran plot scores for all residues in allowed regions are 98.5 %, 98.9% and 99.0% for epac2-cAMP, epac2-BrefA and epac2-BrefA1 respectively (Fig 4 panel e). These results suggest stable confirmation of protein tertiary structure.

## Antibody modelling

### Single domain antibody (sdAb) design

The single domain antibody is labelled as nanobody contain single variable antibody domain. These are designed with antigen specificity and expresses stable interactions.

**Design** was carried out by selecting human variable chain sequence. A best template was 1DEE_B (Fig 5a) with a score of 0.98. The antibody comprises of 118 amino acids and CDR regions of modelled antibody (Fig 5b) are highlighted.

### Benchmarking of Protein-protein docking

Interaction with epac2-Rap1b complex has been studied and validated structure is available is considered as a reference standard. The same two proteins were subjected to Protein-protein docking. The top 10 interaction results showed similar domain/cavity in epac2 protein. 5 out of 9 residues of Rap1b has similar residues interaction with epac2. These results prove the accuracy and efficiency of protein docking protocols. The docking results are shown in (Supplementary Fig 2).

### Epac2 interaction antibody

Two set of antigen-antibody interactions are carried out compare and contrast the efficiency of antibody binding.

### Antibody interaction with simulated epac2

Epac2 protein simulated after interaction with brefA1 is subjected to interaction with Rap1b. The top 10 interactions were analysed and results which showed more stability are compiled in (Supp Table 3). We found that there were no clashes with during the epitope mapping studies. Hydrogen bonding and salt bridge was identified between Arg 100-Asp 331 and Asp72 and Lys 381. Lys 75-Asp 423 showed hydrogen bonding (Supp table 2). The mentioned interactions showed stability proving efficiency. (Fig 5c)

### Mutagenesis of Antibody

From the antibody and simulated epac2 interactions we compiled the most crucial residues of antibody to perform mutagenesis study. The residues S55, Y100A, R100, D72, R19 and K75 were subjected to mutagenesis by their substitution with alternative 19 amino acids to verify if it leads to increase in binding stability and antibody interaction. The results are fluctuations in solvability scores are recorded in (Table 3). It can be observed that R19 substitution with Pro, S55 with Lys, K75 with Pro and Y100A with Pro lead to increase in delta (δ) stability of antibody interactions. A graphical representation of increase in stability for crucial amino acids with individual amino acids in (Fig 5 panel e). These above-mentioned substitutions were performed in the antibody sequence.

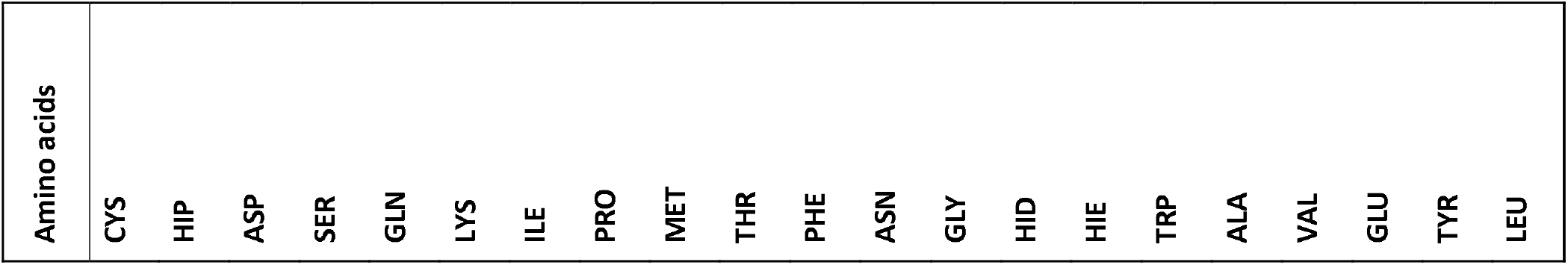

**Table 3.** Mutagenesis table with solvation score.

| TYR:100A | ARG:100 | LYS:75 | ASN:73 | SER:55 | ARG:19 |
| --- | --- | --- | --- | --- | --- |
| 9.25 | 6.35 | 1.82 | 7.38 | 7.69 | 15.43 |
| 3.78 | 0.31 | -9.79 | 7.98 | 16.29 | 8.89 |
| 9.42 | 3.87 | 10.86 | 8.07 | 9.2 | 20.93 |
| 8.84 | 6.23 | 2.6 | 6.92 | 8.12 | 15.32 |
| 6.26 | 1.28 | 4.65 | 0.76 | 0.67 | 13.11 |
| 13.47 | 7.11 | -3.01 | 13.43 | 41.56 | 16.97 |
| 2.57 | 4.94 | 4.63 | -1.6 | 9.72 | 13.72 |
| 38.91 | 26.9 | 56.81 | 6.25 | 34.49 | 32.75 |
| 1.18 | -1.27 | 3.24 | 1.92 | -5.04 | 5.46 |
| 7.73 | 5.15 | 2.37 | 2.34 | 2.84 | 16.57 |
| -1.73 | 0.03 | 2.9 | 3.51 | 6.98 | 16.51 |
| 7.51 | 9.23 | 5.49 | 7.7 | 5.95 | 18.88 |
| 11.28 | 4.41 | -0.31 | 9.4 | 9.41 | 17.83 |
| 1.68 | 1.36 | 1.81 | 3.23 | 9.41 | 12.89 |
| 5.45 | 1.29 | 3.61 | 4.89 | 13.19 | 6.59 |
| 1.67 | -2.42 | -15.75 | 2.18 | 5.91 | 17.45 |
| 3.15 | 4.95 | 5.92 | -1.29 | 38.32 | 16.75 |
| -1.43 | 2.85 | 5.07 | -2.22 | 10.02 | 14.75 |
| 2.3 | -3.23 | 4.36 | 2.92 | 4.86 | 17.76 |
| 10.17 | 1.13 | 2.53 | 0.9 | 12.79 | 15.98 |
| 4.8 | -1.27 | -1.23 | -0.72 | 28.77 | 9.3 |

### Mutated antibody interaction with simulated epac2

The mutated antibody was again subjected to docking with epac2. The results as expected showed increase in interactions with higher number of hydrogen bonding. There was a total of 8 hydrogen bonding interactions and 1 salt bridge (Fig 5d). We also found 2 clashes in the interaction but each followed by a hydrogen bonding each would negate the effects of stearic clashes. The antibody would be bound with increased efficiency. (Supp table 3)

### Antibody interaction with native epac2

A native epac2 is the structural conformation of epac2 before the BrefA1 induced simulation. The results suggest (Supp Table 4) the interaction efficiency to be lower and the docking will not be stable due to presence of clashes. From the top 10 interactions viewed, the best interaction has total of 59 clashes. The interactions also showed formation of 2 salt bridges between Asp 61-Lys 706 and Arg 99-Glu 585. The stability of salt bridges is not efficient enough over the clashes occurring in the interaction. The lack of hydrogen bonding decreases the stability of interactions too. This study provides enough evidence with lack of antibody not interacting with sufficient stable interactions with epac2(Supp table 4).

This study proves that the designed antibody shows preferential binding ability with simulated epac2 over native one. This antibody can be used for identification and quantification of epac2 after the conformational changes with BrefA1.

## Conclusion

The reprofiling of known inhibitor via combinatorial strategy is an efficient protocol to understand the impact of side chain fragments on protein targets. The need to discovery of new compounds can be avoided if the known compounds can be utilized more efficiently via minor substitutions. In the current study, new variants of Brefeldin A were docked against EPAC2 and best docked variant had better binding efficiency than cAMP.

This has a direct impact on the protein conformation, due to the hinge mechanism of EPAC2. The change in conformation of EPAC2 makes its N-terminal domain inaccessible to the RAP1B protein. Herein Brefeldin A, a natural inhibitor due to minor substitution behaves as an antagonist to EPAC2. The known compounds/inhibitors will provide an advantage in its ADME properties over the design of newer compounds. The simulation studies and setting up a minimum time of 200ns will enable better understanding of structural changes on the target due to ligand interactions. The longer interaction period will provide insights into target structure stability which cannot be achieved over smaller simulation runs. This simulation results can avoid minimizing the time and resources needed for *In-vivo* studies by selecting only those interactions which have stable structures by end of simulation course.

*In-silico* design of antibodies prove to be an efficient method in understanding the interaction behaviour. The single domain antibody (sdAb) is specifically designed to bind with EPAC2 protein after binding with BrefA1 and change in conformation. The sdAb has relatively greater binding efficiency over native EPAC2 structure. The study includes CDR grafting and mutagenesis provide more confidence in the results. Single domain antibodies Selectively bind to antigens, less molecular weight, higher heat resistance and stable towards detergents. Diagnostic applications include Fusion with fluorescent protein called chromobody.

Studies including a robust workflow like these can be fruitful by saving time and resource by speeding up the process in wet lab validation.

## Acknowledgements

The authors sincerely thank Schrodinger Inc. for providing software support for method development, conduction and analysis. We thank Dr. Vachaspati Mishra for providing insights into problem statement. Gratitude towards Mr. Vasant Kumar for assistance in developing high resolution images.

## Declaration

### Author Contribution

Methodology, analysis and drafting of manuscript by AUC. VN was involved in refining the methods, conclusions and review of final manuscript. All authors have reviewed the manuscript and agreed for submission.

## Competing interests

The authors claim NO competing interests with any research lab.

## Data Availability

Not applicable

## Funding

No Funding is available for this research

## Supplementary Information

**S1 Supplementary Figure 1: Similarity and sequence coverage of 4F7Z over other EPAC2 crystal structures available in PDB**

The selection of 4F7Z as receptor for docking studies has been justified by providing a comprehensive view of domain and sequence coverage of the selected structure over others. The PDB ids are highlighted in yellow.

**S2 Supplementary table 1: docking score and prime energy of lead like molecules from MM-GBSA output.**

**S3 Supplementary Fig 2: Benchmarking results for Protein-Protein interactions**

The evaluation of protein-protein docking was benchmarked by comparison with known interaction of epac2 with rap1b. The interaction residues are highlighted in red

**S4 Supplementary table 2: Antibody interaction with simulated epac2**

**S5 Supplementary table 3: Mutated antibody interaction with simulated epac2**

**S6 Supplementary table 4: Antibody interaction with native epac2**

## Notes

### Competing Interest Statement

The authors have declared no competing interest.

