## Supplementary figures and images for "Re-profiling of natural inhibitor via combinatorial drug screening: Brefeldin A variant design as an effective antagonist leading to EPAC2 structure modification and antibody design for identification"

### Supplemental Figure 1

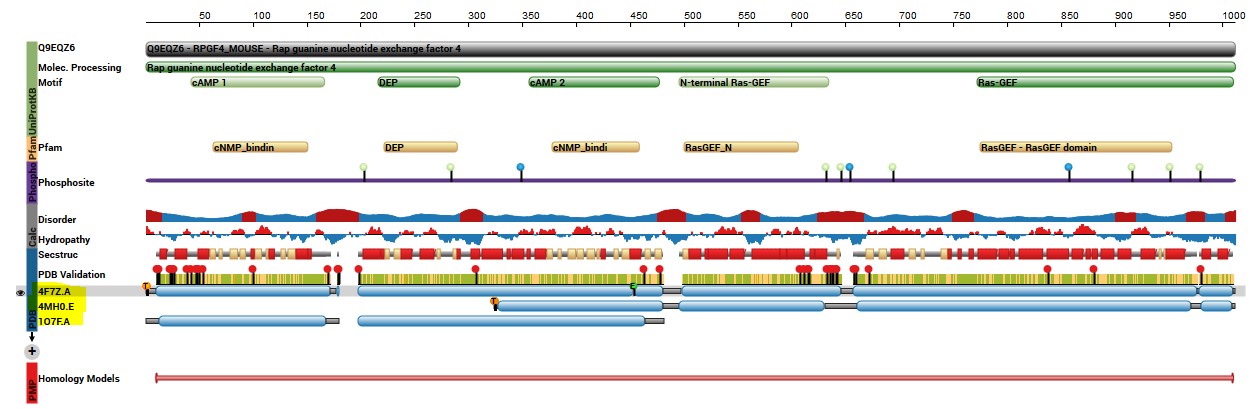

### Supplemental Figure 2

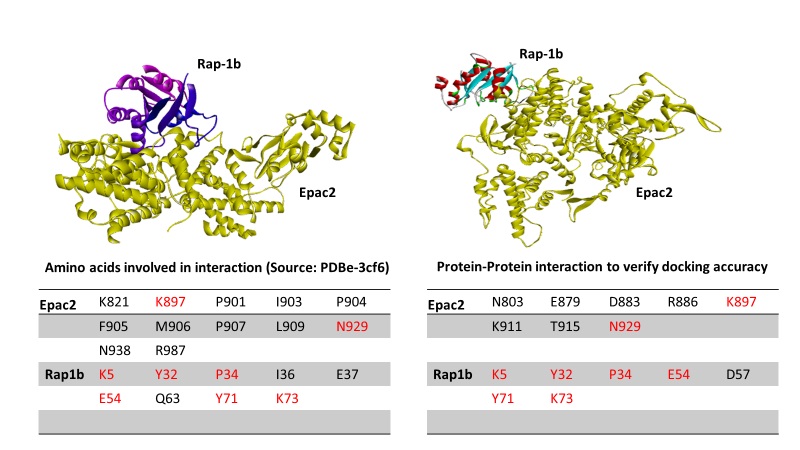
