## Supplemental Table 1 for "Re-profiling of natural inhibitor via combinatorial drug screening: Brefeldin A variant design as an effective antagonist leading to EPAC2 structure modification and antibody design for identification"

| **Title** | **Docking score** | **Energy** |
| --- | --- | --- |
| 5287620; [from N-methylurea 1]; [from imidazolium 2]; [from methanesulfonamide 1] | -10.841 | 37.476 |
| 5287620; [from water 1]; [from imidazolium 2]; [from imidazole 2] | -10.836 | 41.156 |
| 5287620; [from imidazolium 2]; [from methanesulfonamide 1]; [from methanesulfonamide 1] | -10.488 | 24.595 |
| 5287620; [from benzenesulfonamide 1]; [from N-methylacetamide 1]; [from imidazole 2] | -10.056 | 49.575 |
| 5287620; [from ammonium 1]; [from acetamide 3]; [from acetamide 3] | -10.036 | 35.155 |
| 5287620; [from ammonium 1]; [from imidazolium 2]; [from methanesulfonamide 1] | -10.035 | 38.795 |
| 5287620; [from imidazolium 1]; [from water 1]; [from cyclohexane 1] | -10.027 | 41.392 |
| 5287620; [from water 1]; [from imidazolium 1]; [from N-methylacetamide 1] | -10.019 | 48.27 |
| 5287620; [from pyridine 3]; [from acetamide 3]; [from acetamide 3] | -9.989 | 34.233 |
| 5287620; [from imidazolium 2]; [from imidazole 1]; [from water 1] | -9.949 | 45.31 |
| 5287620; [from acetamide 3]; [from piperidinium 1]; [from formate 1] | -9.859 | 41.836 |
| 5287620; [from pyridine 3]; [from acetamide 3]; [from N-methylurea 1] | -9.856 | 35.661 |
| 5287620; [from N-phenylurea 1]; [from imidazolium 2]; [from imidazole 2] | -9.846 | 56.852 |
| 5287620; [from pyridine 2]; [from acetamide 3]; [from methanesulfonamide 1] | -9.837 | 28.119 |
| 5287620; [from imidazolium 1]; [from imidazolium 2]; [from formate 1] | -9.774 | 41.578 |
| 5287620; [from imidazolium 1]; [from imidazolium 2]; [from methanesulfonamide 1] | -9.77 | 39.061 |
| 5287620; [from N-methylacetamide 2]; [from imidazolium 2]; [from imidazole 2] | -9.701 | 41.382 |
| 5287620; [from water 1]; [from acetamide 3]; [from methanesulfonamide 1] | -9.69 | 25.062 |
| 5287620; [from N-methylurea 1]; [from pyridine 1]; [from N-methylmethanesulfonamide 1] | -9.69 | 34.823 |
| 5287620; [from pyridine 3]; [from N-methylurea 1]; [from N-methylurea 1] | -9.684 | 42.157 |
| 5287620; [from N-methylmethanesulfonamide 1]; [from imidazolium 2]; [from acetamide 2] | -9.658 | 30.439 |
| 5287620; [from water 1]; [from imidazolium 2]; [from methanesulfonamide 1] | -9.648 | 32.1 |
| 5287620; [from imidazolium 1]; [from acetamide 1]; [from trifluoromethane 1] | -9.635 | 25.862 |
| 5287620; [from N-phenylurea 1]; [from imidazolium 2]; [from imidazole 1] | -9.627 | 56.308 |
| 5287620; [from water 1]; [from acetamide 3]; [from pyridine 2] | -9.621 | 35.03 |
| 5287620; [from pyridine 2]; [from acetamide 3]; [from N-methylurea 1] | -9.56 | 40.353 |
| 5287620; [from ammonium 1]; [from imidazolium 2]; [from trifluoromethane 1] | -9.533 | 41.476 |
| 5287620; [from imidazolium 1]; [from N-methylurea 1]; [from methanesulfonamide 1] | -9.517 | 39.402 |
| 5287620; [from methanesulfonamide 3]; [from imidazolium 2]; [from methanesulfonamide 1] | -9.506 | 25.238 |
| 5287620; [from water 1]; [from acetamide 3]; [from imidazole 2] | -9.495 | 42.841 |
| 5287620; [from imidazolium 1]; [from methanesulfonamide 3]; [from N-methylmethanesulfonamide 1] | -9.484 | 31.244 |
| 5287620; [from benzene 1]; [from N-methylurea 1]; [from N-methylurea 1] | -9.458 | 51.377 |
| 5287620; [from methanesulfonamide 1]; [from methanesulfonamide 3]; [from N-methylmethanesulfonamide 1] | -9.427 | 10.876 |
| 5287620; [from N-methylurea 1]; [from N-methylurea 1]; [from N-methylmethanesulfonamide 1] | -9.388 | 36.599 |
| 5287620; [from formate 1]; [from imidazole 3]; [from N-methylmethanesulfonamide 1] | -9.352 | 21.115 |
| 5287620; [from benzene 1]; [from acetamide 3]; [from N-methylurea 1] | -9.31 | 44.174 |
| 5287620; [from imidazole 3]; [from imidazolium 2]; [from methanesulfonamide 1] | -9.3 | 34.34 |
| 5287620; [from acetamide 3]; [from acetamide 3]; [from imidazole 2] | -9.251 | 42.61 |
| 5287620; [from N-methylmethanesulfonamide 1]; [from acetamide 3]; [from imidazole 2] | -9.205 | 31.977 |
| 5287620; [from methane 1]; [from N-methylacetamide 1]; [from tetrazolate 1] | -9.189 | 37.012 |
| 5287620; [from acetamide 1]; [from imidazolium 2]; [from formate 1] | -9.179 | 28.931 |
| 5287620; [from imidazolium 1]; [from water 1]; [from imidazole 2] | -9.165 | 42.896 |
| 5287620; [from formate 1]; [from methanesulfonamide 1]; [from imidazole 2] | -9.159 | 27.062 |
| 5287620; [from acetamide 1]; [from N-methylurea 1]; [from N-methylmethanesulfonamide 1] | -9.151 | 25.79 |
| 5287620; [from ammonium 1]; [from acetamide 1]; [from methanesulfonamide 1] | -9.114 | 21.621 |
| 5287620; [from formate 1]; [from N-methylurea 1]; [from imidazole 2] | -9.107 | 45.383 |
| 5287620; [from acetamide 3]; [from imidazolium 1]; [from acetamide 2] | -9.105 | 38.485 |
| 5287620; [from water 1]; [from methanesulfonamide 2]; [from N-methylmethanesulfonamide 1] | -9.102 | 17.152 |
| 5287620; [from imidazole 2]; [from N-phenylurea 1]; [from formate 1] | -9.087 | 57.436 |
| 5287620; [from acetamide 1]; [from acetamide 3]; [from imidazole 2] | -9.083 | 37.596 |
| 5287620; [from acetamide 3]; [from imidazolium 2]; [from acetamide 1] | -9.082 | 32.612 |
| 5287620; [from water 1]; [from N-methylurea 1]; [from methanesulfonamide 1] | -9.078 | 25.732 |
| 5287620; [from ammonium 1]; [from piperidinium 1]; [from imidazole 2] | -9.069 | 56.474 |
| 5287620; [from N-methylurea 1]; [from imidazolium 2]; [from imidazole 3] | -9.065 | 43.398 |
| 5287620; [from methanesulfonamide 1]; [from imidazole 2]; [from methanesulfonamide 1] | -9.064 | 18.693 |
| 5287620; [from N-methylurea 1]; [from imidazolium 2]; [from N-methylmethanesulfonamide 1] | -9.061 | 34.215 |
| 5287620; [from water 1]; [from methane 1]; [from pyridine 2] | -9.056 | 32.157 |
| 5287620; [from acetamide 1]; [from imidazolium 2]; [from acetamide 2] | -9.055 | 32.945 |
| 5287620; [from methanesulfonamide 1]; [from imidazole 2]; [from formate 1] | -9.055 | 31.401 |
| 5287620; [from benzene 1]; [from N-methylurea 1]; [from N-methylacetamide 1] | -9.051 | 57.247 |
| 5287620; [from imidazolium 2]; [from imidazolium 2]; [from tetrazolate 1] | -9.042 | 43.324 |
| 5287620; [from N-phenylurea 1]; [from methanesulfonamide 1]; [from methanesulfonamide 1] | -9.039 | 30.958 |
| 5287620; [from acetamide 2]; [from imidazolium 2]; [from formate 1] | -9.03 | 33.249 |
| 5287620; [from propane 1]; [from imidazolium 2]; [from methanesulfonamide 1] | -9.021 | 33.742 |
| 5287620; [from methanesulfonamide 1]; [from acetamide 3]; [from formate 1] | -9.009 | 26.901 |
| 5287620; [from methanesulfonamide 1]; [from imidazolium 2]; [from methanesulfonamide 3] | -9.008 | 25.295 |
| 5287620; [from imidazolium 1]; [from methanesulfonamide 3]; [from formate 1] | -9.006 | 30.08 |
| 5287620; [from water 1]; [from methanesulfonamide 1]; [from acetamide 1] | -9.005 | 21.191 |
| 5287620; [from water 1]; [from N-methylurea 1]; [from pyridine 2] | -9.003 | 41.407 |
| 5287620; [from water 1]; [from imidazole 2]; [from N-methylmethanesulfonamide 1] | -9.001 | 23.67 |
| 5287620; [from N-methylurea 1]; [from imidazolium 2]; [from N-methylmethanesulfonamide 2] | -8.961 | 39.674 |
| 5287620; [from methane 1]; [from water 1]; [from N-methylmethanesulfonamide 1] | -8.961 | 19.089 |
| 5287620; [from imidazole 1]; [from imidazolium 2]; [from imidazole 2] | -8.958 | 46.695 |
| 5287620; [from water 1]; [from imidazolium 2]; [from hydrogen cyanide 1] | -8.947 | 34.003 |
| 5287620; [from acetamide 1]; [from water 1]; [from cyclohexane 1] | -8.946 | 28.151 |
| 5287620; [from acetamide 1]; [from hydrogen cyanide 1]; [from imidazole 2] | -8.945 | 32.078 |
| 5287620; [from N-methylacetamide 1]; [from pyridine 1]; [from N-methylacetamide 1] | -8.938 | 59.324 |
| 5287620; [from water 1]; [from imidazolium 1]; [from methane 1] | -8.933 | 35.35 |
| 5287620; [from imidazolium 2]; [from imidazolium 2]; [from methane 1] | -8.926 | 47.982 |
