## Supplemental Table 2 for "Re-profiling of natural inhibitor via combinatorial drug screening: Brefeldin A variant design as an effective antagonist leading to EPAC2 structure modification and antibody design for identification"

| **Residue** | **Closest** | **Distance** | **Specific Interactions** | **# HB** | **# Salt Bridges** | **# Pi Stacking** | **# Disulfides** | **# vdW Clash** | **Surface Complementarity** | **Buried SASA** |
| --- | --- | --- | --- | --- | --- | --- | --- | --- | --- | --- |
| E:310:Met | H:55:Ser | 4.0 A |  | 0 | 0 | 0 | 0 | 0 | 0.71 | 47.10% |
| E:311:Arg |  |  |  | 0 | 0 | 0 | 0 | 0 | 0 | 30.60% |
| E:312:Met |  |  |  | 0 | 0 | 0 | 0 | 0 | 0 | 6.30% |
| E:331:Asp |  |  |  | 0 | 0 | 0 | 0 | 0 | 0 | 17.90% |
| E:335:His |  |  |  | 0 | 0 | 0 | 0 | 0 | 0 | 34.40% |
| E:381:Lys | H:72:Asp | 3.2 A | 1x hb, 1x salt bridge to H:72:Asp | 1 | 1 | 0 | 0 | 0 | 0.64 | 53.70% |
| E:398:His |  |  |  | 0 | 0 | 0 | 0 | 0 | 0 | 26.50% |
| E:399:Glu | H:19:Arg | 3.6 A |  | 0 | 0 | 0 | 0 | 0 | 0 | 43.20% |
| E:422:Glu |  |  |  | 0 | 0 | 0 | 0 | 0 | 0.28 | 22.60% |
| E:423:Asp | H:75:Lys | 3.2 A | 1x hb to H:75:Lys | 1 | 0 | 0 | 0 | 0 | 0.22 | 19.60% |
| E:424:Asn |  |  |  | 0 | 0 | 0 | 0 | 0 | 0.13 | 14.60% |
| E:426:His |  |  |  | 0 | 0 | 0 | 0 | 0 | 0 | 0.60% |
