## Supplemental Table 3 for "Re-profiling of natural inhibitor via combinatorial drug screening: Brefeldin A variant design as an effective antagonist leading to EPAC2 structure modification and antibody design for identification"

| Residue | Closest | Distance | Specific Interactions | # HB | # Salt Bridges | # Pi Stacking | # Disulfides | # vdW Clash | Surface Complementarity | Buried SASA |
| --- | --- | --- | --- | --- | --- | --- | --- | --- | --- | --- |
| E:448:Arg | H:100:Arg | 3.6 A |  | 0 | 0 | 0 | 0 | 0 | 0.45 | 18.50% |
| E:455:Asp | H:100:Arg | 2.9 A | 2x hb to H:100:Arg | 2 | 0 | 0 | 0 | 0 | 0.69 | 44.00% |
| E:456:Val | H:100A:Tyr H:100:Arg | 3.5 A 3.9 A |  | 0 | 0 | 0 | 0 | 0 | 0.62 | 24.20% |
| E:457:Leu | H:100A:Tyr | 3.4 A |  | 0 | 0 | 0 | 0 | 0 | 0.52 | 100.00% |
| E:458:Val | H:100A:Tyr | 3.5 A |  | 0 | 0 | 0 | 0 | 0 | 0.26 | 99.60% |
| E:482:Val |  |  |  | 0 | 0 | 0 | 0 | 0 | 0 | 5.10% |
| E:483:Met | H:100C:Tyr H:99:Arg | 2.9 A 3.5 A |  | 0 | 0 | 0 | 0 | 0 | 0.93 | 65.20% |
| E:484:Ser | H:100A:Tyr H:99:Arg H:100C:Tyr | 3.0 A 3.2 A 3.2 A | 1x hb to H:99:Arg 1x clash to H:100A:Tyr | 1 | 0 | 0 | 0 | 1 | 0.91 | 100.00% |
| E:485:Gly | H:100C:Tyr H:100A:Tyr | 3.1 A 3.4 A |  | 0 | 0 | 0 | 0 | 0 | 0.86 | 100.00% |
| E:486:Thr | H:100A:Tyr | 3.4 A |  | 0 | 0 | 0 | 0 | 0 | 0.83 | 53.90% |
| E:487:Pro |  |  |  | 0 | 0 | 0 | 0 | 0 | 0.33 | 39.10% |
| E:490:Ile |  |  |  | 0 | 0 | 0 | 0 | 0 | 0.49 | 40.20% |
| E:513:Asn |  |  |  | 0 | 0 | 0 | 0 | 0 | 0.4 | 19.30% |
| E:514:Asp | H:100E:Tyr | 3.0 A | 1x hb to H:100E:Tyr | 1 | 0 | 0 | 0 | 0 | 0.74 | 29.20% |
| E:515:Phe | H:100E:Tyr | 3.7 A |  | 0 | 0 | 0 | 0 | 0 | 0.58 | 65.90% |
| E:516:Val |  |  |  | 0 | 0 | 0 | 0 | 0 | 0.26 | 100.00% |
| E:517:Met | H:103:Trp H:37:Val H:45:Leu | 3.3 A 3.3 A 3.4 A |  | 0 | 0 | 0 | 0 | 0 | 0.83 | 84.70% |
| E:518:Met | H:93:Ala H:100E:Tyr H:103:Trp H:100F:Gly H:100D:Ala H:94:Lys | 3.1 A 3.2 A 3.3 A 3.3 A 3.6 A 3.9 A |  | 0 | 0 | 0 | 0 | 0 | 0.83 | 99.90% |
| E:519:His |  |  |  | 0 | 0 | 0 | 0 | 0 | 0.55 | 100.00% |
| E:520:Cys | H:47:Trp | 3.2 A |  | 0 | 0 | 0 | 0 | 0 | 0.6 | 100.00% |
| E:521:Val | H:100D:Ala H:35:Hie H:47:Trp H:50:Thr | 2.8 A 3.3 A 3.5 A 3.8 A |  | 0 | 0 | 0 | 0 | 0 | 0.86 | 80.20% |
| E:522:Phe | H:100D:Ala H:100E:Tyr | 3.2 A 3.7 A |  | 0 | 0 | 0 | 0 | 0 | 0.66 | 60.80% |
| E:524:Pro | H:58:Tyr | 3.5 A |  | 0 | 0 | 0 | 0 | 0 | 0.67 | 77.10% |
| E:525:Asn |  |  |  | 0 | 0 | 0 | 0 | 0 | 0.13 | 81.50% |
| E:526:Thr |  |  |  | 0 | 0 | 0 | 0 | 0 | 0 | 24.10% |
| E:568:Ala |  |  |  | 0 | 0 | 0 | 0 | 0 | 0 | 4.00% |
| E:569:Met | H:45:Leu H:44:Gly | 3.0 A 3.2 A | 1x hb to H:45:Leu | 1 | 0 | 0 | 0 | 0 | 0.64 | 49.70% |
| E:570:Tyr | H:44:Gly | 3.5 A |  | 0 | 0 | 0 | 0 | 0 | 0.5 | 95.60% |
| E:571:Gly |  |  |  | 0 | 0 | 0 | 0 | 0 | 0 | 20.80% |
| E:572:Asp | H:43:Lys H:40:Ala | 3.3 A 3.9 A |  | 0 | 0 | 0 | 0 | 0 | 0.72 | 49.10% |
| E:573:Leu | H:46:Glu H:44:Gly H:45:Leu | 3.3 A 3.3 A 3.5 A |  | 0 | 0 | 0 | 0 | 0 | 0.86 | 97.90% |
| E:576:Glu | H:62:Ser H:38:Arg H:46:Glu | 2.8 A 2.9 A 3.3 A | 1x hb, 1x salt bridge, 1x clash to H:38:Arg 1x hb to H:62:Ser | 2 | 1 | 0 | 0 | 1 | 0.71 | 81.90% |
| E:577:Asp | H:61:Asp H:60:Thr | 3.3 A 3.3 A | 1x clash to H:60:Thr 1x hb to H:61:Asp | 1 | 0 | 0 | 0 | 1 | 0.59 | 92.80% |
