## Supplemental table 4 for "Re-profiling of natural inhibitor via combinatorial drug screening: Brefeldin A variant design as an effective antagonist leading to EPAC2 structure modification and antibody design for identification"

| Residue | Closest | Distance | Specific Interactions | # HB | # Salt Bridges | # Pi Stacking | # Disulfides | # vdW Clash | Surface Complementarity | Buried SASA |
| --- | --- | --- | --- | --- | --- | --- | --- | --- | --- | --- |
| A:524:Pro |  |  |  | 0 | 0 | 0 | 0 | 0 | 0 | 8.30% |
| A:525:Asn |  |  |  | 0 | 0 | 0 | 0 | 0 | 0 | 1.70% |
| A:526:Thr | H:100E:Tyr | 0.9 A | 17x clash to H:100E:Tyr | 0 | 0 | 0 | 0 | 17 | 0.13 | 64.10% |
| A:575:Gln | H:100A:Tyr H:100C:Tyr | 2.3 A 3.5 A | 1x clash to H:100A:Tyr | 0 | 0 | 0 | 0 | 1 | 0.3 | 76.30% |
| A:576:Glu | H:100A:Tyr H:100C:Tyr | 2.5 A 3.2 A | 1x clash to H:100A:Tyr | 0 | 0 | 0 | 0 | 1 | 0.76 | 56.80% |
| A:577:Asp |  |  |  | 0 | 0 | 0 | 0 | 0 | 0.15 | 48.10% |
| A:578:Asp | H:100C:Tyr H:96:Pro | 0.7 A 2.3 A | 3x clash to H:96:Pro 24x clash to H:100C:Tyr | 0 | 0 | 0 | 0 | 27 | 0.1 | 98.00% |
| A:579:Val | H:100E:Tyr H:100F:Gly | 2.8 A 3.9 A | 8x clash to H:100E:Tyr | 0 | 0 | 0 | 0 | 8 | 0.39 | 75.60% |
| A:581:Met | H:100C:Tyr | 2.6 A | 3x clash to H:100C:Tyr | 0 | 0 | 0 | 0 | 3 | 0.4 | 100.00% |
| A:582:Ala |  |  |  | 0 | 0 | 0 | 0 | 0 | 0.07 | 27.00% |
| A:585:Glu | H:99:Arg | 1.9 A | 1x salt bridge, 4x clash to H:99:Arg | 0 | 1 | 0 | 0 | 4 | 0.21 | 43.00% |
| A:612:Val |  |  |  | 0 | 0 | 0 | 0 | 0 | 0.42 | 48.30% |
| A:613:Lys |  |  |  | 0 | 0 | 0 | 0 | 0 | 0 | 0.30% |
| A:615:Ile | H:100:Arg | 3.8 A |  | 0 | 0 | 0 | 0 | 0 | 0.77 | 9.30% |
| A:616:Ser | H:99:Arg H:100:Arg | 2.4 A 3.1 A | 4x clash to H:99:Arg 1x clash to H:100:Arg | 0 | 0 | 0 | 0 | 5 | 0.49 | 98.60% |
| A:617:Glu |  |  |  | 0 | 0 | 0 | 0 | 0 | 0 | 4.20% |
| A:618:Asp | H:100:Arg | 1.6 A | 10x clash to H:100:Arg | 0 | 0 | 0 | 0 | 10 | 0.28 | 26.90% |
| A:619:Ala | H:100:Arg | 3.5 A |  | 0 | 0 | 0 | 0 | 0 | 0 | 8.10% |
| A:620:Lys | H:100:Arg | 2.9 A |  | 0 | 0 | 0 | 0 | 0 | 0.9 | 5.50% |
| A:621:Ala | H:100:Arg | 2.9 A | 1x clash to H:100:Arg | 0 | 0 | 0 | 0 | 1 | 0.79 | 60.50% |
| A:649:Ser | H:100A:Tyr | 1.6 A | 20x clash to H:100A:Tyr | 0 | 0 | 0 | 0 | 20 | 0.31 | 96.60% |
| A:650:Asp |  |  |  | 0 | 0 | 0 | 0 | 0 | 0.62 | 22.80% |
| A:651:Glu |  |  |  | 0 | 0 | 0 | 0 | 0 | 0 | 6.20% |
| A:669:Pro |  |  |  | 0 | 0 | 0 | 0 | 0 | 0 | 34.80% |
| A:673:Ser |  |  |  | 0 | 0 | 0 | 0 | 0 | 0 | 7.70% |
| A:675:Lys | H:58:Tyr H:57:Lys H:59:His | 1.5 A 3.7 A 3.9 A | 39x clash to H:58:Tyr | 0 | 0 | 0 | 0 | 39 | 0.19 | 65.10% |
| A:676:Glu |  |  |  | 0 | 0 | 0 | 0 | 0 | 0 | 12.50% |
| A:706:Lys | H:61:Asp | 3.4 A | 1x salt bridge, 1x clash to H:61:Asp | 0 | 1 | 0 | 0 | 1 | 0.87 | 25.90% |
| A:707:Ser | H:58:Tyr | 2.7 A | 2x clash to H:58:Tyr | 0 | 0 | 0 | 0 | 2 | 0.72 | 59.70% |
| A:708:Asn | H:58:Tyr | 3.6 A |  | 0 | 0 | 0 | 0 | 0 | 0.48 | 36.80% |
| A:789:Lys |  |  |  | 0 | 0 | 0 | 0 | 0 | 0 | 46.90% |
